# Cross-regulation of viral kinases with cyclin A secures shutoff of host DNA synthesis

**DOI:** 10.1101/856435

**Authors:** Boris Bogdanow, Max Schmidt, Henry Weisbach, Iris Gruska, Barbara Vetter, Koshi Imami, Eleonore Ostermann, Wolfram Brune, Matthias Selbach, Christian Hagemeier, Lüder Wiebusch

**Affiliations:** Labor für Pädiatrische Molekularbiologie, Charité Universitätsmedizin Berlin, Augustenburger Platz 1, 13353 Berlin, Germany; Research group „Proteome Dynamics“, Max Delbrück Center for Molecular Medicine, Robert-Rössle-Str. 10, 13125 Berlin, Germany; Research group „Structural Interactomics“, Leibniz Forschungsinstitut für Molekulare Pharmakologie, Robert-Rössle-Str. 10, 13125 Berlin, Germany; Haupt Pharma Berlin GmbH, Moosrosenstr. 7, 12347 Berlin, Germany; Laboratory of Molecular & Cellular BioAnalysis, Kyoto University, 606-8501 Kyoto, Japan; Heinrich Pette Institute, Leibniz Institute for Experimental Virology, Martinistr. 52, 20251 Hamburg, Germany; Charité-Universitätsmedizin Berlin, Charitéplatz 1, 10117 Berlin, Germany

**Author notes:** These authors contributed equally (Boris Bogdanow, Max Schmidt).

## Abstract

Herpesviruses encode conserved protein kinases to stimulate phosphorylation-sensitive processes during infection. How these kinases bind to cellular factors and how this impacts their regulatory functions is poorly understood. Here, we use quantitative proteomics to determine cellular interaction partners of human herpesvirus (HHV) kinases. We find that these kinases can target key regulators of transcription and replication. The interaction with Cyclin A and associated factors is identified as a specific signature of β-herpesvirus kinases. Cyclin A is recruited via RXL-motifs that overlap with nuclear localization signals (NLS) and locate in the non-catalytic N-terminal regions. This architecture is conserved for viral kinases of HHV6, HHV7 and rodent CMVs. Docking to Cyclin A competes with NLS function, enabling dynamic changes in kinase localization and substrate phosphorylation. The viral kinase redirects Cyclin A to the cytosol, which is essential for the inhibition of cellular DNA replication during infection. Our data highlight a fine-tuned and physiologically important interplay between a cellular cyclin and viral kinases.

## INTRODUCTION

Herpesviruses are a wide-spread family of large, double-stranded DNA viruses replicating within the nuclei of their host cells (Pellett & Roizman, 2013). Generally, herpesviruses are highly adapted to their hosts and persist in a latent mode of infection unless immunodeficiency provokes viral reactivation and disease (Adler et al., 2017). They have diversified into three subfamilies: neurotropic α-herpesviruses, broadly infective β-herpesviruses and lymphotropic γ-herpesviruses (Davison, 2010). Despite their widely different pathogenic properties and clinical manifestations, all herpesviruses share a common set of conserved core genes (Mocarski, 2011), mostly encoding essential structural components and replication factors. Among the few core genes with regulatory function are the conserved herpesvirus protein kinases (CHPKs) (Gershburg and Pagano, 2008), which are medically important both as drug targets (Li and Hayward, 2013) and as prodrug activating enzymes (Topalis et al., 2018).

CHPKs belong to the group of viral serine/threonine kinases (Jacob et al., 2011). Although possessing a considerable sequence divergence, CHPKs have a number of common characteristics including auto-phosphorylation, nuclear localization, incorporation into the tegument layer of virus particles and phosphorylation of other tegument proteins (Jacob et al., 2011). CHPKs target regulators of the DNA damage checkpoint (Li et al., 2011; Tarakanova et al., 2007), phosphorylate the translational elongation factor EF-1δ (Kato et al., 2001) and counteract the IRF3-dependent type I interferon response (Hwang et al., 2009; Wang et al., 2009).

CHPKs were reported to preferentially target cellular cyclin-dependent kinase (CDK) phosphorylation sites (Kawaguchi et al., 2003; Oberstein et al., 2015). In particular, the kinases of human β- and γ-herpesviruses show significant structural and functional homology to CDKs (Hamirally et al., 2009; Kuny et al., 2010; Lee et al., 2007; Romaker et al., 2006). This has led to their designation as viral CDK-like kinases (v-CDKs) (Kuny et al., 2010). V-CDKs lack amino acids that are known to be essential for interaction of cellular CDKs with cyclins, CDK inhibitors (CKIs) and CDK activating kinases (CAKs). Accordingly, v-CDKs are considered immune to cellular control mechanisms (Hume et al., 2008), despite the fact that the UL97 kinase of human cytomegalovirus (human herpesvirus 5, HHV5) can physically interact with various cyclins (Graf et al., 2013; Steingruber et al., 2019). V-CDKs mimic CDK1 and 2 in phosphorylating Lamin A/C (Hamirally et al., 2009; Kuny et al., 2010), retinoblastoma-associated tumor suppressor RB1 (Hume et al., 2008; Kuny et al., 2010; Prichard et al., 2008) and deoxynucleoside triphosphate (dNTP) hydrolase SAMHD1 (Businger et al., 2019; Kim et al., 2019; Zhang et al., 2019). Collectively, the CDK-like activities of CHPKs facilitate the nuclear egress of virus capsids (Milbradt et al., 2016) and secure the supply of dNTPs and cellular replication factors for viral DNA replication (Deutschmann et al., 2019; Zhang et al., 2019).

Significant knowledge has accumulated over the recent years about CHPK functions. It is known that CHPKs can recruit cellular proteins as substrates via conserved docking motifs (Iwahori et al., 2015; Prichard et al., 2008). Further, CHPKs can be regulated by their interaction partners and even exert kinase independent functions (Avey et al., 2016; Li et al., 2012). However, a systematic analysis of CHPK protein-protein interactions is lacking and data linking interactions to functions of the kinase are scarce. To broaden our understanding of CHPKs, we used an affinity-purification mass-spectrometry (AP-MS) approach and identified shared and differential interaction partners of CHPKs from all herpesvirus subfamilies. We found that Cyclin A-CDK complexes build a common set of interactors for β-herpesvirus v-CDKs. Taking mouse cytomegalovirus (MCMV, murid herpesvirus 1) as a model system, we show that during infection the stoichiometric formation of cytoplasmic v-CDK and Cyclin A assemblies causes a global shift in substrate phosphorylation and the viral shutoff of host DNA synthesis. Collectively, our data demonstrates that herpesvirus kinases have evolved as protein-interaction hubs that can recruit a rich repertoire of cellular proteins. The functional versatility of β-herpesvirus v-CDKs is underpinned by their ability to crosstalk with a cellular cyclin via direct, non-catalytic interaction.

## RESULTS

### Conserved HHV kinases target key regulators of transcription and replication

Systems-level approaches have provided important insights into the molecular functions of CHPKs. For example, yeast 2 hybrid screens revealed several binary binding partners of CHPKs (Calderwood et al., 2007; Uetz et al., 2006) and phosphoproteome profiling was successfully used to assess CHPK substrate specificity (Li et al., 2011, 2015; Oberstein et al., 2015; Umaña et al., 2018). However, comprehensive and comparative information about CHPK interaction partners at the proteome level is lacking.

To identify CHPK interaction partners, we transfected SILAC (stable isotope labeling of amino acids in cell culture) heavy and light labeled HEK293-T cells with HA-tagged CHPKs of seven different human herpesviruses (HHVs) or vector controls. We performed these experiments in triplicates, including label-swaps (Figure 1A), and subjected the samples to HA affinity purification and shotgun proteomics. We observed an overall good reproducibility of individual replicates (Figure S1A). To discriminate candidate interactors from background binders, they were required to fall below a p-value of 0.05 and meet a SILAC fold-change cut off at 1 % false discovery rate (FDR). (Figures 1B, S1B-C). In total, we quantified 2,816 proteins, of which 135 were classified as candidate interaction partners to at least one kinase (Table S1). This list includes several previously found CHPK interactors and substrates, such as SAMHD1 (Zhang et al., 2019), IRS4, CCAR2 (Steingruber et al., 2016), RBL2 (Iwahori et al., 2017), PPP2CA, PUM1, PRDX1 (Li et al., 2011) and NUMA1, TUBA1B, MAP7D1(Umaña et al., 2018).

**Figure 1.**
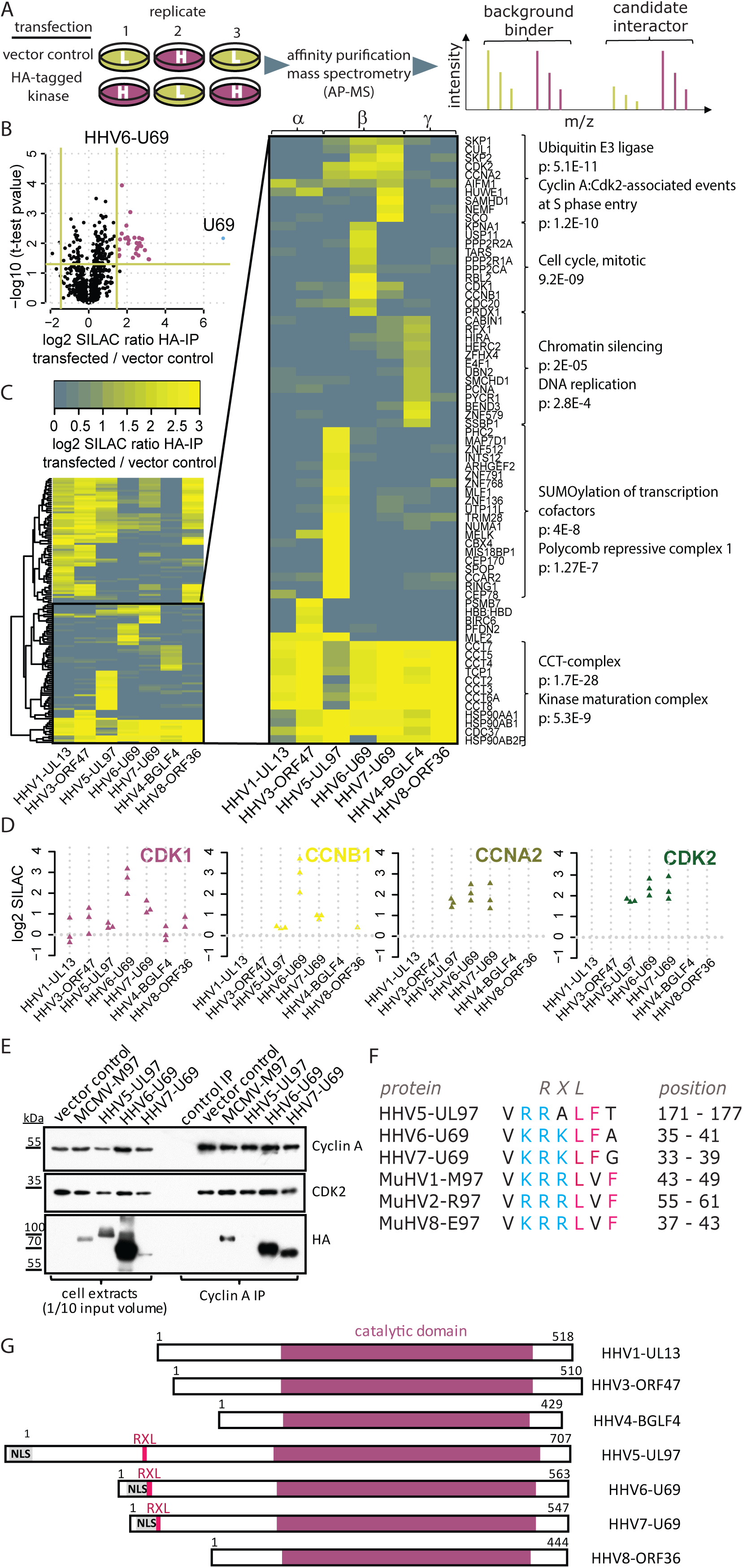
Conserved HHV kinases target key regulators of cellular replication and transcription. **(A)** Experimental setup. SILAC heavy and light labeled HEK293-T cells were transfected with constructs encoding HA-tagged versions of human herpesviral kinases or control vectors in triplicates. Samples were subjected to anti-HA affinity purification followed by mass-spectrometry. **(B)** Candidate selection. Candidate interaction partners were discriminated from background binders based on a combination of the t-test p-value of three replicates and the mean of the SILAC ratios. P-value cut-off at 0.05 and fold-change cut-off at FDR=1%. A volcano plot for the HHV6-U69 kinase is exemplarily shown. **(C)** Cross-comparison of kinase interaction partners. All candidate interaction partners were compared for co-enrichment across the indicated kinases of α-, β- and γ-HHVs. Enrichment profiles were clustered and selected sets of clusters were subjected to GO-enrichment (inlet and brackets). **(D)** Cyclin-CDK complexes co-precipitate with β-herpesviral kinases. The SILAC ratios for individual replicates for the prey proteins CDK1, CCNB1, CCNA2 and CDK2 across AP-MS samples for the indicated kinases. **(E)** HEK293-T cells were transfected with expression vectors encoding HA-tagged versions of the indicated kinases or an empty control vector. Whole cell extracts were prepared at 2 days post transfection and subjected to Cyclin A co-IP. The output material was analyzed by immunoblotting for the presence of Cyclin A, CDK2 and HA-tagged kinases. **(F)** Alignment of putative Cyclin A-binding sites in human and rodent β-herpesvirus kinases. Basic residues are highlighted in blue, bulky hydrophobic residues in pink. **(G)** Schematic of human herpesviral kinases. Shown are the location of the conserved catalytic domains and the relative positions of RXL/Cy motifs (pink) and predicted or experimentally validated nuclear localization signals (NLS) (grey) within the less-conserved N-terminal parts of the proteins.

We next aimed to determine interaction partners, that were common to all kinases, class-specific (i.e. restricted to either kinases of α, β or γ-classes) or unique to individual kinases. Therefore, we clustered the SILAC fold-changes of candidate interactors across all tested kinases (see Figure 1C). We found the chaperonin containing TCP1 (CCT) and the kinase maturation complex (CDC37, HSP90) to copurify with all kinases analyzed. This confirms the previous observation thatCHPKs interact with the same set of cellular proteins that assist in folding and maturation of cellular kinases (Sun et al., 2013; Taipale et al., 2012).

Proteins that co-precipitated with BGLF4 (HHV4, Epstein-Barr-virus), but were not significantly enriched with other kinases, include factors involved in DNA replication (HERC2, E4F1, PCNA, SSBP1) and chromatin silencing (SMCHD1, CABIN1, HIRA, BEND3). Proteins that precipitated with pUL97 (HHV5) but not other kinases were functionally related to regulation of transcription (PHC2, RING1, CBX4, TRIM28, ZNF136, ZNF791). The cell cycle-related phosphatase PP2A (PPP2CA, PPP2R1A, PPP2R2A), regulators of p53/p21 ubiquitylation (HUWE1, USP11) and nuclear import/export mediators (KPNA1, NEMF) specifically co-enriched with U69 kinases of HHV6 and HHV7.

Importantly, we found that all human β-herpesvirus kinases precipitated S/G2 phase-specific cyclins (Cyclin A / CCNA2, Cyclin B / CCNB1), CDKs (CDK1, CDK2) and associated proteins (SKP1, SKP2). While enrichment of Cyclin A-CDK2 was reproducibly strong, Cyclin B-CDK1 did not enrich with pUL97 in our transfection experiment (Figure 1D). We were able to validate the interaction with Cyclin A-CDK2 by reverse co-immunoprecipitations (co-IPs). Also, we observed this type of interaction for the homologous M97 kinase of mouse cytomegalovirus / murid herpesvirus 1 (MCMV / MuHV-1) (Figure 1E). Collectively our interactome of human CHPKs provides a rich resource and suggests that these kinases are crucially involved in the regulation of transcription, epigenetic remodeling and cell cycle control. Cyclin A-CDK2 complexes build a common subset of interactors for β-herpesvirus kinases, suggesting important functional implications.

### β-herpesvirus kinases interact with cyclin-CDK complexes via NLS-RXL modules

β-herpesvirus kinases lack most of the residues of CDKs that directly interact with cyclins, including a conserved PSTAIRE helix (Hume et al., 2008). Instead, we found RXL/Cy motifs in the N-terminal, non-catalytic parts of β-herpesvirus kinases (Figure 1F-G). Such motifs are typically used for substrate and inhibitor recruitment to cyclin-CDKs (Schulman et al., 1998). Importantly, the positions of the RXL-type sequences within the largely divergent N-termini are well conserved (Figure S2). These putative Cyclin A binding elements in β-herpesvirus kinases of the roseolovirus (HHV6, HHV7) and muromegalovirus (MuHV1, MuHV2, MuHV8) genera are in close proximity to clusters of positively charged residues (Figure S2), which are predicted, and in the case of HHV6 validated, classical bipartite nuclear localization signals (NLS) (Isegawa et al., 2008). In fact, the C-terminal part of the NLS sequences directly overlaps with the N-terminal part of the RXL motifs (Figure 2A). Thus, when we set out to test the contribution of RXL-motifs in U69 and M97 to Cyclin A binding, we had to consider the possibility that RXL mutations may negatively affect NLS function. We therefore designed two mutant versions of each kinase: one disrupting the core of the RXL motif (RXL->AXA) and one changing only the hydrophobic part (LF->AA, LXF->AXA), leaving the basic residues of the NLS intact (Figure 2A). Both mutations abolished binding of M97 and U69 kinases to Cyclin A (Figure 2B). Thus, RXL sequence motifs found in β-herpesvirus kinases act as Cyclin A docking sites. RXL mutation not only prevents Cyclin A binding but also interaction with other cyclin-associated factors found in the interactome analysis (Figure 1C), as exemplarily shown for HHV6-U69 (Figure S3). This indicates that those factors are not direct v-CDK interactors but instead co-recruited with Cyclin A. Thus, the RXL-motif triggers the formation of higher order v-CDK-cyclin-CDK complexes.

**Figure 2.**
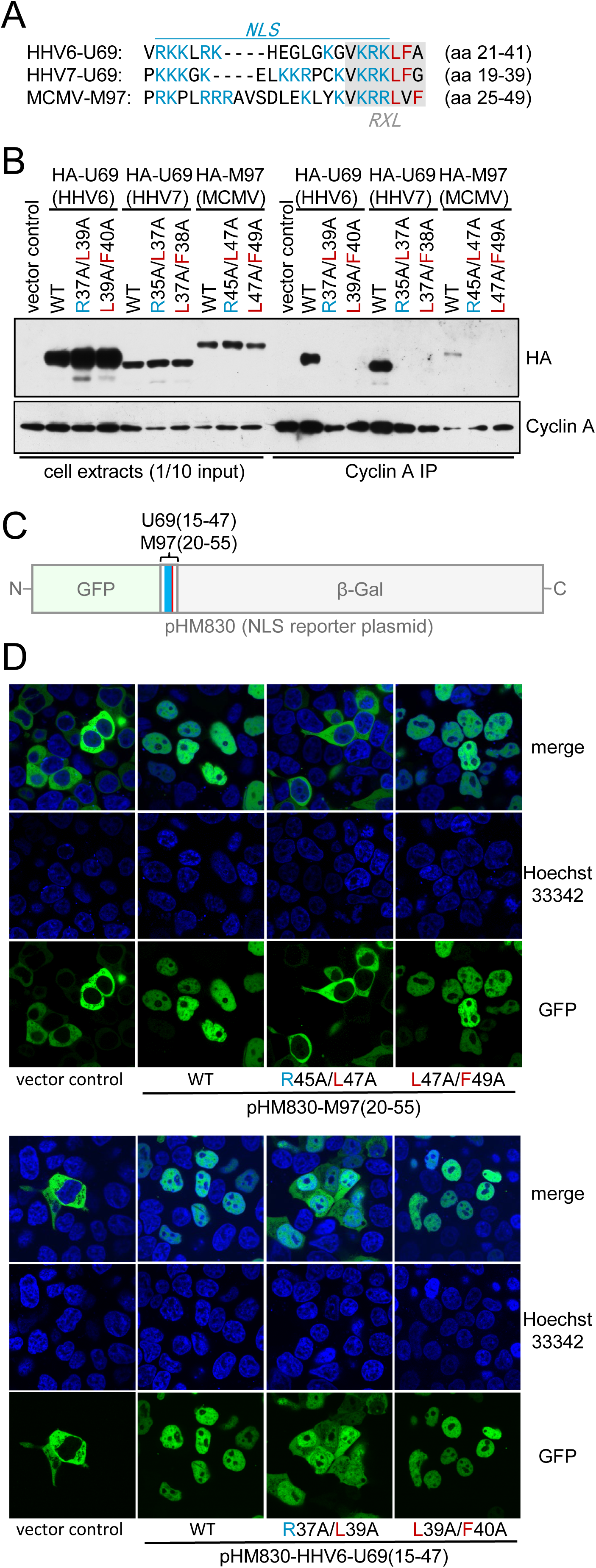
NLS and Cyclin A docking motifs overlap in U69 and M97 kinases. **(A)** Sequence alignment of overlapping NLS-RXL elements in M97 and U69 kinases. Stretches of basic residues characteristic for nuclear localization signals (NLS) are highlighted in blue, conserved leucine and phenylalanine residues participating only in the RXL/Cy motif in pink. A mutation strategy was designed to separate NLS function from cyclin binding. RXL AXA -mutations affect both RXL and NLS sequences, LXF AXA and LF AA mutations only the RXL/Cy motif. **(B)** HEK293-T cells were transfected with expression vectors encoding HA-tagged wild-type (WT) or mutant forms of U69 and M97 kinases. Cyclin A co-IP was carried out at 2 days post transfection. Input material and precipitates were analyzed by anti-HA and anti-Cyclin A immunoblotting for the presence of WT and mutant kinases as well as Cyclin A. **(C)** Sequence fragments encompassing the NLS-RXL region of U69/M97 mutant and wildtype kinases were cloned between and in-frame the green fluorescent protein (GFP) and β-galactosidase genes in pHM830. **(D)** HEK293-T cells were transfected with WT or RXL/Cy-mutant versions of the M97-NLS reporter constructs, as indicated. The subcellular localization of GFP was analyzed at 24 h post transfection using confocal live-cell imaging microscopy. Nuclei were counterstained with Hoechst-33342.

We then assessed the consequences of RXL mutations for NLS function. To this end, we cloned 32-37 amino acid segments encompassing the overlapping NLS and RXL elements of M97 and U69 kinases into an NLS reporter construct (Sorg and Stamminger, 1999) (Figure 2C). Integration of wild type (WT) NLS-RXL regions into the chimeric reporter induced nuclear accumulation of the otherwise cytoplasmic GFP signal (Figure 2D), confirming the predicted NLS activity. RXL to AXA mutations weakened this activity in HHV6-U69 and even disrupted it in M97. In contrast, the NLS function remained intact when only the hydrophobic residues of the RXL/Cy motifs are mutated (Figure 2D). These results demonstrate that cyclin binding and NLS sequences are integrated into a composite motif that can be functionally separated by mutations.

### The viral kinase assembles higher-order cyclin-CDK complexes in infected cells

We then aimed to analyze the NLS-RXL module in a representative infection system and chose MCMV for ease of manipulation. We first assessed the time resolved protein interactome of M97 using SILAC and AP-MS (Figure S4A, Table S2). Early during infection (12 h), the M97 interactome consisted almost exclusively of cyclins, CDKs and associated proteins (Figure S4B). In the late phase (36 h), additional viral and cellular factors co-purified with M97 (Figure S4D), including M50/M53, the nuclear egress complex of MCMV (Muranyi et al., 2002) and Lin54, the DNA-binding subunit of the cell cycle-regulatory MuvB complex (Marceau et al., 2016). Taken together, this indicates that M97 functions in cell cycle regulation, viral egress and control of gene expression.

In consistence with the data from transfected cells, M97 interacted with Cyclin A/B-CDK complexes throughout infection (Figure 3A, Figure S4B,D). In particular, Cyclin A-CDK2 was present in similar molar amounts as M97 itself in HA-M97 precipitates (Figure S4C,E) suggesting a strong and stoichiometric interaction in infected cells. Mutation of the RXL/Cy motif (Figure S5) disrupted the interaction of Cyclin A with M97 (Figure 3C-D). However, these mutations did not influence Cyclin A protein levels (Figure 3B) or Cyclin A-associated kinase activity (Figure 3F). Moreover, RXL/Cy mutations, in contrast to the the “kinase dead” K290Q mutation (Deutschmann et al., 2019), did not compromise M97 levels and activity (Figure 3B,E). Thus, M97-Cyclin A binding has no influence on abundance and enzymatic activity of the involved interaction partners.

**Figure 3.**
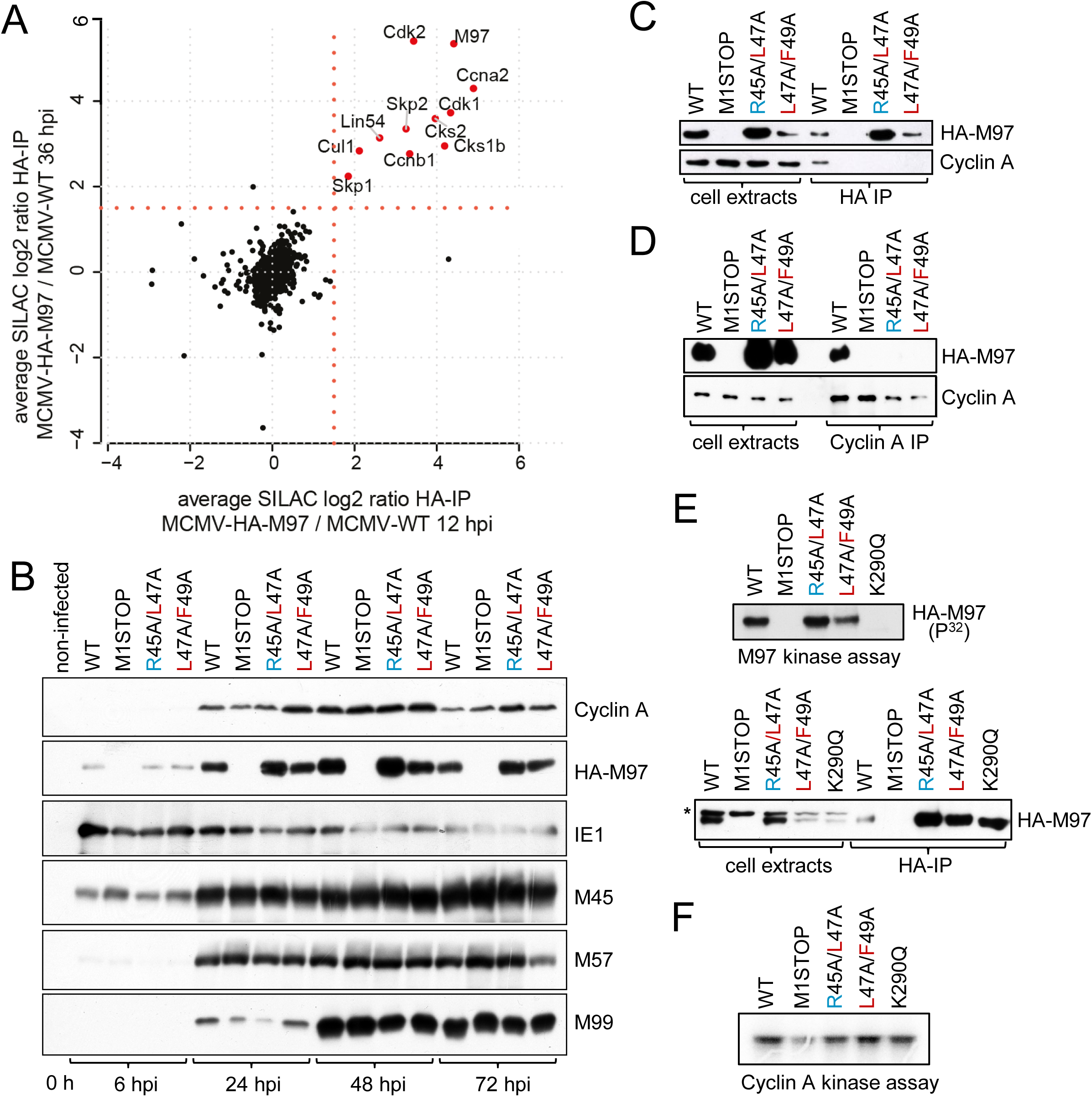
Formation of higher-order cyclin-CDK assemblies with a viral kinase in infected cells. **(A)** A time-resolved interactome of M97 during MCMV infection (see also Figure S4). Enrichment ratios for proteins co-precipitating with HA-M97 at 12 or 36 hours post infection in a scatterplot. The cut-off at SILAC fold-change 1.5 is indicated by a red dotted line. **(B-F)** Serum-starved 3T3 fibroblasts were infected with the indicated MCMV-HA-M97 variants. **(B)** The expression of Cyclin A and of selected immediate-early, early and late MCMV gene products was monitored by immunoblot analysis. **(C-D)** Immunoprecipitation of the indicated HA-M97 variants by anti-HA-coupled paramagnetic microbeads **(C)** or reverse immunoprecipitation **(D)** confirms RXL/Cy-dependent interaction with Cyclin A-CDK2. **(E-F)** Assays for M97 **(E)** and cyclin A **(F)** enzymatic activity. Lysates from MCMV-infected 3T3 fibroblasts were subjected to anti-HA-IP **(E)** or cyclin A-IP **(F)** and used as input material for γ-P32-ATP kinase assay. Recombinant Histone H1 was used as RXL/Cy-independent Cyclin A substrate and autophosphorylation of M97 was measured for the M97 kinase assay. Incorporation of γ-P32-ATP was visualized by autoradiography.

### Cyclin binding controls the subcellular localization of M97

In respect of the overlapping NLS and RXL/C motifs within M97, we next tested whether Cyclin A binding alters the subcellular localization of M97. To address this, we chose a system where Cyclin A levels are dynamically changing. In quiescent cells, Cyclin A is absent at very early time points of infection (Figure 3B) and induced between 6 and 24 h by MCMV (Figure 3B). Using immunofluorescence microscopy, we observed that M97-WT is nuclear at 6 h post infection (Figure 4A), when Cyclin A is still low, but redistributes to a predominantly cytoplasmic localization pattern at 24 and 48 h post infection (see Figure 4A, middle left column), when Cyclin A levels are high. Thus, the appearance of M97 in the cytosol correlates with increased Cyclin A expression.

**Figure 4.**
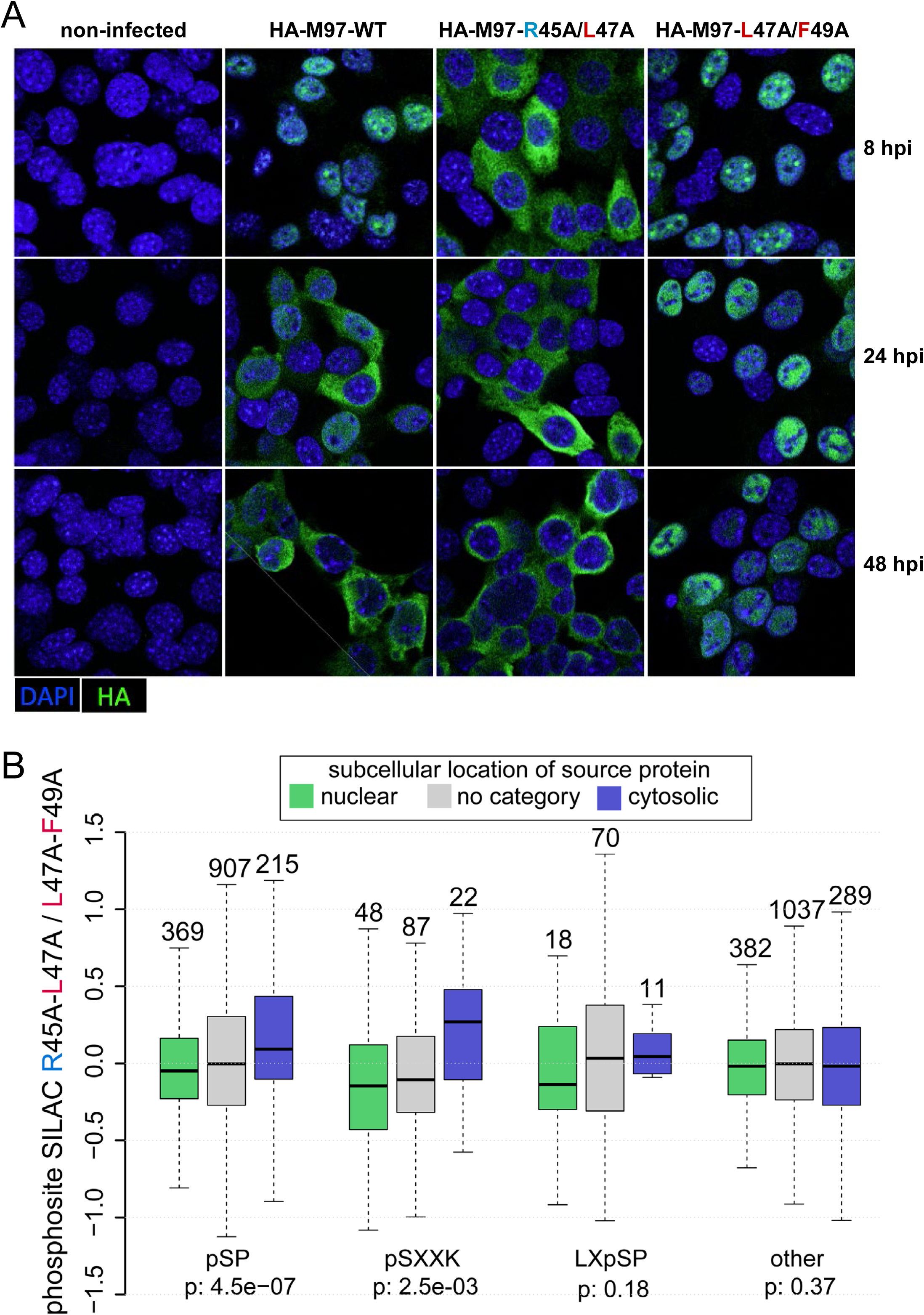
The RXL-NLS module controls the subcellular localization of M97 and the phosphorylation status of its substrates. **(A)** Serum starved 3T3 fibroblasts were infected with the indicated MCMV-HA-M97 variants or left uninfected. At 8 h, 24 h and 48 h post infection, cells were examined by confocal immunofluorescence microscopy for subcellular localization of HA-M97 (green). Nuclei were counterstained with DAPI. Representative images are shown. **(B)** SILAC labeled 3T3 cells were serum starved and infected with NLS-RXL/Cy-deficient M97^R45A/L47A^ or RXL/Cy-deficient M97^L47A/F49A^ viruses (see also Figure S6). At 24 h post infection cells were harvested and subjected to a phospho-proteomic workflow. Boxplots for phospho-sites that belong to proteins with nuclear, cytoplasmic or uncategorized subcellular GO annotation are depicted with their SILAC log2 fold-change between M97^R45A/L47A^ and M97^L47A/F49A^ infection. pSP, pSXXK, LXpSP or all other sites were assessed individually. Data-points outside the whiskers were omitted in the figure. P-values are based on one sided wilcoxon rank-sum test comparing nuclear and cytosolic subsets.

Then we tested the contribution of the RXL/NLS module to the time dependent change in M97 localization. When the NLS function was disrupted by R45A/L47A mutation, M97 was confined to the cytoplasm throughout infection (see Figure 4A, middle right column). In contrast, when the NLS part of the NLS-RXL module was left intact and only Cyclin binding was prevented (L47A/F49A), M97 showed a constitutively nuclear localization pattern (Figure 3A, right column). Thus, Cyclin A binding is responsible for the late relocalization of M97. This provides evidence for a scenario, where Cyclin A masks the NLS and thereby effectively interferes with nuclear entry of M97.

Next, we studied how the dynamic and Cyclin A-dependent relocalization of M97 impacts protein phosphorylation. Therefore, we compared cells that were either infected with RXL deficient MCMV-M97^L47A/F49A^ or NLS-RXL deficient MCMV-M97^R45A/L47A^. The two mutant viruses reflect the early nuclear or late cytosolic localization of M97 during MCMV-WT infections. We globally assessed substrate phosphorylation by performing a proteomic analysis of phosphopeptides and whole cell lysates of SILAC-labeled cells at 24 h post infection (Figure S6A, Table S3).

First, we corrected phospho-site ratios for protein level changes (Wu et al., 2011) (Figure S6B-C). Next, we categorized proteins and corresponding phosphosites based on their GO annotation as nuclear, cytoplasmic or unclear (“no category”) (Figure S6D). Then, we specifically analyzed phospho-sites that match known v-CDK target motifs (Oberstein et al., 2015), such as pS/TP, pSXXK, LXpSP (p denotes the phosphorylated residue). When cells were infected with M97^R45A/L47A^ mutant, pS/TP and pSXXK sites residing in the cytosol were significantly stronger phosphorylated than nuclear sites (Figure 4B). In contrast, when cells were infected with M97^L47A/F49A^ mutant, the same set of sites were stronger phosphorylated when they belonged to nuclear proteins. The most pronounced differences in the target spectrum were observed when phospho-serines followed by prolines were positioned between hydrophobic amino acids and lysines (Figure S6E). Collectively, these data argue that Cyclin A binding enables switch-like changes in the substrate spectrum and subcellular distribution of M97.

### M97 controls cell cycle progression by cytoplasmic sequestration of Cyclin A

The binding of M97 to Cyclin A may have functional consequences not only for M97 but also for Cyclin A. Cyclin A is essential for cellular DNA replication and cell cycle progression from S phase to mitosis (Girard et al., 1991). Importantly, Cyclin A function depends on its nuclear localization (Cardoso et al., 1993; Maridor et al., 1993). Therefore, we analyzed whether M97 binding impacts the subcellular distribution of Cyclin A. We found Cyclin A, and to a lesser extent CDK2, to be depleted from the nucleus in MCMV-WT infected cells (see Figure 5A). This effect was Cyclin A-specific as Cyclin E was evenly distributed between nuclear and cytoplasmic fractions. Mutation of the M97 translation start site prevented the cytoplasmic enrichment of Cyclin A. Mutations of the RXL/Cy motif in M97 even led to a predominant nuclear localization of Cyclin A. Thus, Cyclin A is sequestered within the cytosol dependent on its binding to M97.

**Figure 5.**
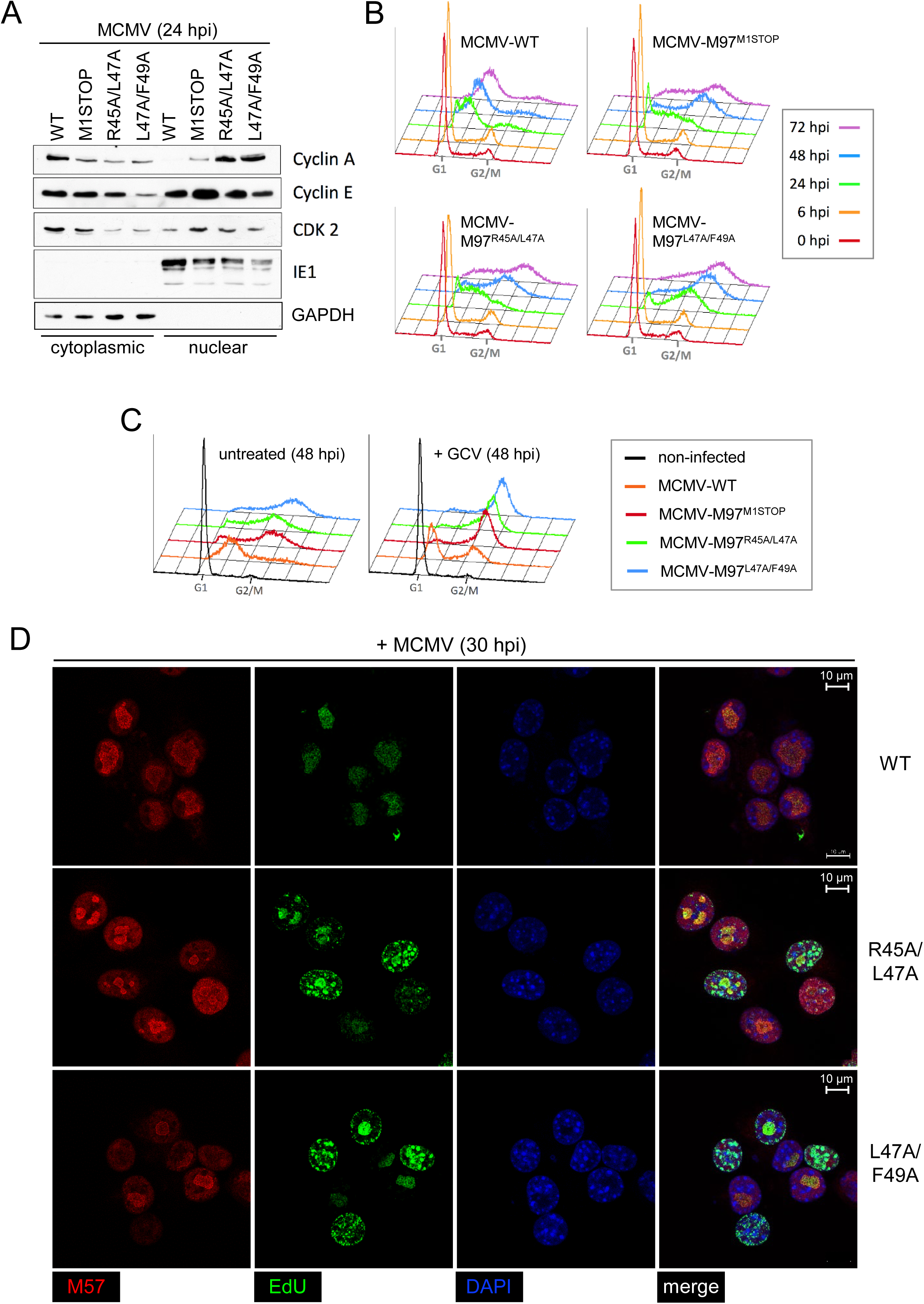
Cytosolic sequestration of Cyclin A by M97 is essential for viral shutoff of host DNA synthesis. Serum-starved 3T3 cells were infected with the indicated recombinant viruses and subjected to subcellular fractionation **(A)**, cell cycle analysis **(B,C)** or confocal microscopy **(D). (A)** The levels of Cyclin A, E and Cdk2 proteins was determined at 24 h post infection in nuclear and cytosolic fractions by immunoblotting. The soluble viral nuclear protein IE1 and the cytosolic marker GAPDH served as controls. **(B, C)** The DNA content of infected cells was analyzed by propidium iodide staining followed by flow cytometry and plotted as DNA histograms. **(B)** The accumulation of viral and cellular DNA was monitored over the time course of infection. **(C)** To discriminate viral from cellular DNA replication, infected cells were treated with ganciclovir (GCV) or left untreated. **(D)** At 30 hpi, infected cells were pulse-labeled with EdU. EdU staining via click-chemistry (green fluorescence) served to determine sites of cellular and viral DNA synthesis. EdU was combined with immunofluorescence detection of the viral replication factor M57 (red fluorescence) that marks sites of viral DNA synthesis. DAPI was used for nuclear counterstaining (blue fluorescence). Representative images are shown.

This observation prompted us to investigate whether the nuclear depletion of Cyclin A by M97 affects cell cycle progression. Therefore, we measured the DNA content of quiescent fibroblasts infected with WT and M97 mutant viruses by flow cytometry (Figure 5B). We found a ganciclovir-sensitive increase of viral DNA in MCMV-WT infected cells, consistent with previous reports (Wiebusch et al., 2008). In contrast, M97 mutant viruses caused a rapid and ganciclovir-resistant accumulation of cells with a G2/M DNA content. This shows that loss of M97 expression or M97-Cyclin A interaction leads to S phase entry and cell cycle progression to G2/M phase. Very similar results were obtained in primary, non-immortalized cells (MEFs), the only difference being that here the RXL/Cy mutation leads to a cellular DNA content that exceeds the normal 4n DNA content of diploid G2/M cells (Figure S7). This indicates that the M97 mutant virus bypasses cellular control mechanisms protecting from DNA over-replication.

We then aimed to confirm the crucial role of M97-Cyclin A interaction for inhibition of cellular DNA synthesis by an orthogonal approach. We chose to combine EdU pulse labeling and fluorescence microscopy to spatially discriminate sites of active viral or cellular DNA synthesis. In MCMV-WT infected cells, DNA synthesis is confined to nuclear replication compartments that stain positive for M57 (Figure 5D), a viral ssDNA binding protein (Strang et al., 2012). In contrast, cells infected with M97-RXL/Cy mutants incorporated EdU in foci distributed over the whole nucleus, indicating that the restriction of cellular DNA replication was lost. The M97 mutant infected cells also showed a reduced size of viral replication compartments, indicating less efficient viral DNA synthesis. Thus, cytoplasmic sequestration of M97-Cyclin A complexes serves as an essential viral mechanism to shut off competing host DNA synthesis during productive infection.

## DISCUSSION

Here we present the first comparative analysis of CHPK interactomes. Our analysis led to the identification of host-protein interaction signatures that are common, class-specific or unique to CHPKs (Figure 1C). The interaction of β-herpesvirus-CHPKs with Cyclin A orchestrates higher-order complex formation of cell cycle regulatory factors (Figure 3). Notably, the interaction is regulatory and has functional consequences for both, the CHPK and Cyclin A. For MCMV, it results in the dynamic relocalization of the viral kinase (Figure 4A) and consequently an altered substrate spectrum (Figure 4B). Further, the interaction leads to cytosolic sequestration of Cyclin A, which is essential for viral arrest of DNA replication (Figure 5).

V-CDKs are a subset of CHPKs that share key aspects of host CDK-function and the ability to complement for CDK activity in yeast cells (Hamirally et al., 2009; Hume et al., 2008; Kuny et al., 2010). V-CDKs have lost sequence features enabling control by cellular factors, such as CAK, CKI and cyclins (Hume et al., 2008). Instead, we show that some v-CDKs acquired RXL motifs (Figure 1), which are typically used by cellular CDK substrates and inhibitors for recognition by cyclins (Adams et al., 1996; Schulman et al., 1998). These motifs enable a regulatory cross-talk of Cyclin A and v-CDKs, which is characterized by cell cycle dependent regulation of the viral kinase on the one hand (Figure 4) and neutralization of Cyclin A in stoichiometric protein assemblies on the other hand (Figure S4, Figure 5). The latter effect of v-CDK-Cyclin A interaction is reminiscent of CKI function (Besson et al., 2008). Thus, we propose that β-herpesviruses integrate the antipodal activities of CDKs and CKIs on one gene product. This combination allows a β-herpesvirus kinase like M97 to activate S/G2 metabolism (Deutschmann et al., 2019) while inhibiting cellular DNA synthesis (Figure 5).

An exception among human β-herpesviruses is HCMV, which has evolved a different gene product for neutralization of Cyclin A. HCMV produces the small protein pUL21a, which targets Cyclin A for proteasomal degradation (Caffarelli et al., 2013; Eifler et al., 2014). Therefore, it seems that HCMV has shifted its Cyclin A-antagonistic, CKI function from its kinase to pUL21a. In that context it is interesting that the RXL motif in HCMV-pUL97 does not overlap with an NLS (Webel et al., 2012), suggesting an alternative function of pUL97-Cyclin A interaction (Figure 1).

Short linear motifs (SLiMs) can be rapidly acquired by viruses and other pathogens to target host proteins (Chemes et al., 2015; Davey et al., 2011; Via et al., 2015). Here we found that within HHV6, HHV7 and rodent CMV kinases two such SLiMs are fused into one regulatory sequence element (Figure 2). The physical overlap of RXL/Cy and NLS motifs is facilitated by their sequence composition as both contain contiguous stretches of basic amino acids. NLS motifs function as docking sites for nuclear import factors, mainly importin-α (Lange et al., 2007). Accordingly, a docking competition mechanism makes binding to Cyclin A and nuclear localization of M97 mutually exclusive, enabling switch-like changes in viral kinase function (Figure 6A). This puts β-herpesvirus kinases in a row with a number of cellular and viral proteins known to control nucleo-cytoplasmic localization via intermolecular NLS-masking (Christie et al., 2016), with NF-κB as the best understood example (Huxford et al., 1998; Jacobs and Harrison, 1998).

**Figure 6.**
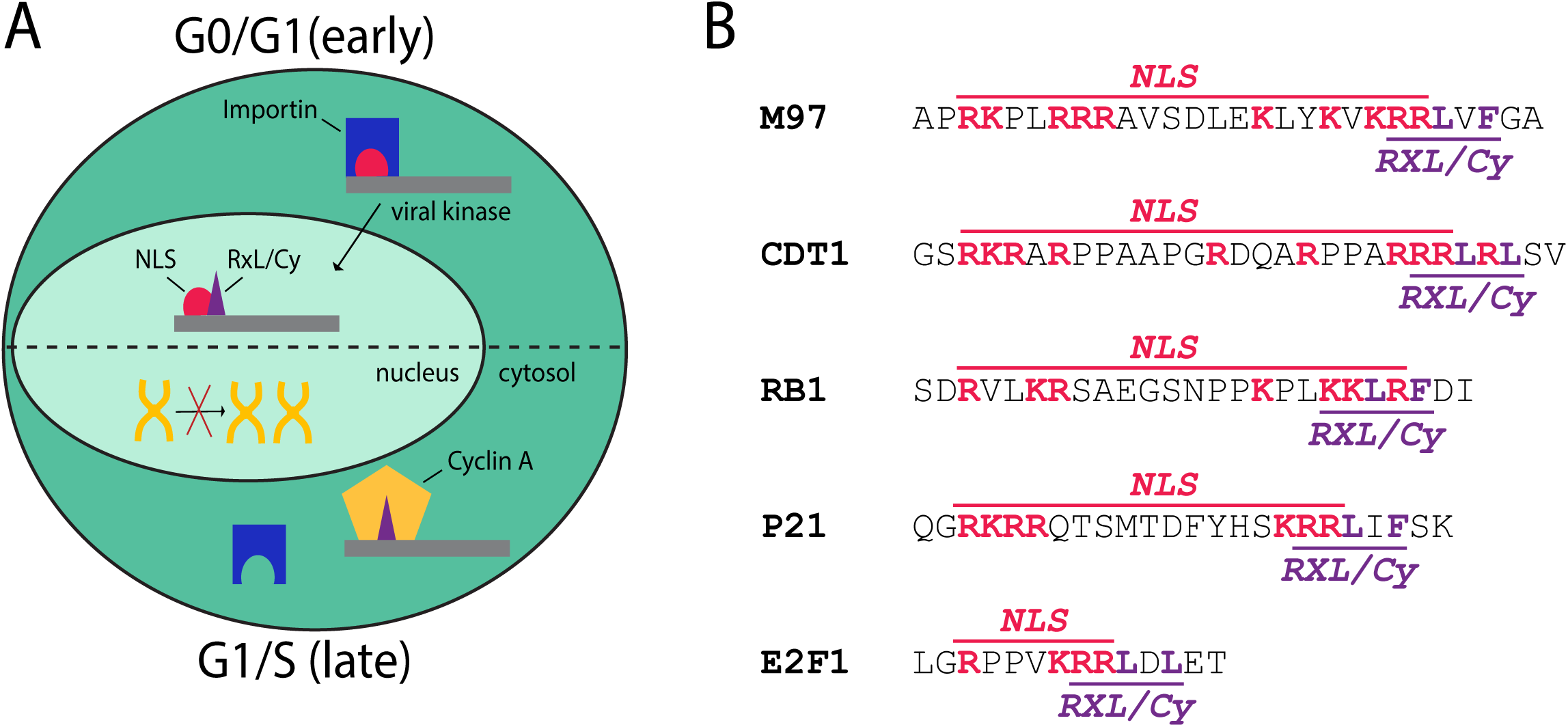
The bi-functional NLS-RXL/Cy module is conserved across several cell cycle regulatory proteins. **(A)** A model summarizing the function of the NLS-RXL/Cy module in infected cells. In the absence of cyclin A (G0/G1, early), the NLS of M97 is functional and M97 is imported into the nucleus. MCMV induces cyclin A and drives the cell cycle towards an S-phase environment (G1/S, late). Cyclin A binds to the RXL/Cy motif on M97 and masks the NLS, leading to cytosolic accumulation of M97-cyclin A complexes. Cellular DNA synthesis is inhibited due to mislocalized Cyclin A. **(B)** Conservation of RXL/Cy-NLS modules across several cellular cell cycle regulatory proteins.

Remarkably, composite NLS-RXL/Cy elements are apparent in a number of key regulatory proteins of the host cell cycle. Although independently described, NLS and RXL/Cy motifs are overlapping in RB1 (Adams et al., 1999; Zacksenhaus et al., 1993), p21/CDKN1A (Adams et al., 1996; Chen et al., 1996; Rodríguez-Vilarrupla et al., 2002), E2F1 (Krek et al., 1994; Müller et al., 1997) and CDT1 (Liu et al., 2004; Nishitani et al., 2004; Sugimoto et al., 2004) (Figure 6B). For these cellular factors, it could be important to consider a docking competition between Cyclin A and importins, a possibility which is so far unexplored. Specifically, cyclin binding may be essential for the cell cycle dependent localization of these proteins (Coqueret, 2003; Jiao et al., 2008; Müller et al., 1997; Tanaka and Diffley, 2002). In other cases, such as CDC6, RXL motifs are adjacent to but not overlapping with NLS. There, Cyclin A recruits CDKs to neutralize the NLS by phosphorylation (Delmolino et al., 2001; Petersen et al., 1999).

In addition to the functionally important interaction of β-herpesvirus kinases with Cyclin/CDKs, our unbiased proteomic survey indicates that CHPKs interact with many more cellular proteins (Figure 1C, Supplementary Table S1). Consistent with previous reports, many CHPK interaction partners are functionally related to DNA repair (Li et al., 2011) and cell cycle control (Kuny et al., 2010). Remarkably, we found that CHPKs also interact with a variety of prominent transcriptional repressor complexes. For example, pUL97 interacts with KAP1/TRIM28-ZNF complexes, known to silence CMV gene expression in stem cells depending on the phosphorylation status of TRIM28 (Rauwel et al., 2015). In addition, pUL97 precipitates with members of the polycomb repressive complex 1 (PRC1; PHC2, RING1, CBX4), which was linked to epigenetic control of herpesvirus gene expression (Hopcraft et al., 2018). Likewise, we found BGLF4 to precipitate with core subunits of the HIRA histone chaperone complex (HIRA, CABIN1 and UBN2). This complex restricts lytic infection by depositing histone H3.3 on incoming herpesviral genomes (Rai et al., 2017). Collectively, these data indicate that CHPKs can target host-derived master regulators of viral transcription, governing the decision between lytic and latent infection programs. This could allow tegument-delivered CHPKs to actively influence the outcome of herpesvirus infections.

Our interactome analysis indicates that CHPKs target master regulators involved in host replication, DNA repair and transcription. It is important to note that this type of experiment cannot resolve protein interactions that depend on the environment of an infected cell. It is thus critical to interpret our findings in the broader context of kinase-associated functions that dynamically change during infection. For instance, phosphorylation of the nuclear protein SAMHD1 could be maintained by the M97 kinase early during infection (Deutschmann et al., 2019) when M97 localizes nuclear (Figure 4A). During later stages of infection, the nuclear lamina is locally disassembled and egress of viral capsids to the cytoplasm occurs. The shift from nuclear replication to cytosolic assembly is accompanied by the concurrent relocalization of the viral kinase. Thus, Cyclin A-interaction helps the kinase to exert infection-stage specific functions.

## MATERIALS AND METHODS

### Cells

HEK293-T cells and NIH-3T3 fibroblasts were cultivated in Dulbecco’s modified Eagle medium (DMEM) supplemented with 10% fetal (293T) or newborn (3T3) bovine serum, 2 mM L-alanyl-L-glutamine, 100 U/ml penicillin, and 100µg/ml streptomycin. Where indicated, cells were synchronized in G0/G1 phase by 48 h growth factor deprivation (0.05% serum). In preparation for proteomic analysis, cells were SILAC-labeled for at least 5 passages using lysine and arginine-deprived DMEM, supplemented with 10% dialyzed serum (cutoff: 10 kDa), 200 mg/l L-proline (only cells destined for phosphoproteome analysis) to minimize arginine to proline conversion (Bendall et al., 2008), heavy (L-[^13^C_6_,^15^N_2_]-lysine (Lys8), L-[^13^C_6_,^15^N_4_]-arginine (Arg10)), medium (L-[^2^H_4_]-lysine (Lys4), L-[^13^C_6_]-arginine (Arg6)) or light (natural lysine (Lys0) and arginine (Arg0)) amino acids. Labeling efficiency and arginine-proline conversion was checked using LC-MS/MS.

### Viruses

Viruses were derived from the m129-repaired MCMV strain Smith bacterial artificial chromosome (BAC) pSM3fr-MCK-2fl (Jordan et al., 2011). Infections were carried out at 37°C under conditions of centrifugal enhancement. In brief, after a virus adsorption period of 30 min, cell cultures were centrifuged for 30 min at 1000 g. Then, the virus inoculum was replaced by fresh medium. Virus titers were determined by flow cytometry of IE1 fluorescent cells at 6 h post infection. Unless otherwise stated, a multiplicity of infection (MOI) of 5 IE protein forming units (IU) per cell was used for experiments.

### Bacmids

MCMV-HA-M97 and MCMV-M97-K290Q were described previously (Deutschmann et al., 2019). R45A/L47A, L47A/F49A and M1STOP mutations were introduced into MCMV-HA-M97 by traceless BAC mutagenesis (Tischer et al., 2010). The oligonucleotide primers used for BAC mutagenesis are specified in Table S4. All mutants were controlled by diagnostic PCR and sequencing (Figure S5). To reconstitute infectious virus, BACs together with pp71 expression plasmid were transfected into 3T3 fibroblasts using an Amaxa nucleofector (Lonza).

### Plasmids

PCGN-based expression plasmids for HA-tagged HHV1-UL13, HHV3-ORF47, HHV4-BGLF4, HHV5-UL97, HHV6-U69, HHV7-U69 and HHV8-ORF36 (Addgene plasmids #26687, #26689, #26691, #26693, #26695, #26697, #26698) were gifts from Robert Kalejta (Kuny et al., 2010). PCIneo-3HA-M97, the expression plasmid for HA-tagged M97, was described previously (Deutschmann et al., 2019). RXL/Cy mutations were introduced by site-directed inverse PCR-mutagenesis (primers see Table S4). PHM830 (Addgene plasmid #20702) was a gift from Thomas Stamminger (Sorg and Stamminger, 1999). Fragments encompassing the NLS-RXL/Cy modules of M97 and U69 were PCR-amplified and cloned between NheI and XbaI sites of pHM830 (primers listed in Table 4). Plasmids were confirmed by Sanger sequencing and purified by cesium chloride/ethidium bromide equilibrium centrifugation. PEI MAX (Polysciences), transfection grade linear polyethylenimine hydrochloride with a molecular weight of 40 kDa, was used for transfection (Longo et al., 2013).

### Phosphopeptide enrichment

SILAC-labeled cells were harvested 24 hours post MCMV infection via scraping. Cells were lysed with 6M urea/2M thiourea in 0.1 M Tris-HCl, pH 8.0. Samples were reduced with 10 mM DTT and alkylated with 50 mM iodoacetamide for 30 min in the dark. Proteins were digested by lysyl endopeptidase (Wako Pure Chemicals) at an enzyme-to-protein ratio of 1:100 (w/w) for 3 h. Subsequently, samples were diluted with 50 mM ammonium bicarbonate to a final concentration of 2 M urea. Digestion with proteomics-grade modified trypsin (Promega) was performed at a enzyme-to-protein ratio of 1:100 (w/w) under constant agitation for 16 h. Enzyme activity was quenched by acidification with trifluoroacetic acid (TFA). The peptides were desalted with C18 Stage Tips (Rappsilber et al., 2003) prior to nanoLC-MS/MS analysis (whole cell lysate samples). An aliquot of the whole cell lysates were further processed for phosphopeptide enrichment. The tryptic digests corresponding to 200 μg protein per condition were desalted with big C18 Stage Tips packed with 10 mg of ReproSil-Pur 120 C18-AQ 5-μm resin (Dr. Maisch GmbH). Peptides were eluted with 200 μL loading buffer, consisting of 80% acetonitrile (ACN) and 6% TFA (vol/vol). Phosphopeptides were enriched using a microcolumn tip packed with 0.5 mg of TiO_2_ (Titansphere, GL Sciences). The TiO_2_ tips were equilibrated with 20 μL of loading buffer via centrifugation at 100 g. 50 μL of each sample were loaded on a TiO_2_ tip via centrifugation at 100 g and this step was repeated until all the sample was loaded. The TiO_2_ column was washed with 20 μL of the loading buffer, followed by 20 μL of 50% ACN/0.1% TFA (vol/vol)). The bound phosphopeptides were eluted using successive elution with 30 μL of 5% ammonium hydroxide and 30 μL of 5% piperidine (Kyono et al., 2008). Each fraction was collected into a fresh tube containing 30 μL of 20% formic acid. 3 μL of 100% formic acid was added to further acidify the samples. The phosphopeptides were desalted with C18 Stage Tips prior to nanoLC-MS/MS analysis.

### Affinity purification

At 12 and 36 hours post infection (M97 interactome) or 1 day post transfection (HHV kinases interactomes), cells were harvested by scraping in PBS. After centrifugation at 300 g, cell pellets were lysed in 25 mM Tris-HCl (pH 7.4), 125 mM NaCl, 1 mM MgCl_2_, 1% Nonidet P-40 (NP-40), 0.1% SDS, 5% glycerol, 1 mM dithiothreitol (DTT), 2 µg/ml aprotinin, 10 µg/ml leupeptin, 1 µM pepstatin, 0.1 mM Pefabloc. For HA affinity purification, a µMACS HA isolation kit (Miltenyi Biotec) was employed according to the manufacturer’s instructions, with the following modifications. The first washing step was carried out using lysis buffer. Lysis buffer without detergent was used for the following washing step. The final washing buffer was 25 mM Tris-HCl (pH 7.4). Samples were eluted in a total volume of 0.2 ml 8 M guanidine hydrochloride at 95°C. Proteins were precipitated from the eluates by adding 1.8 ml LiChrosolv ethanol and 1 µl GlycoBlue. After incubation at 4°C overnight, samples were centrifuged for 1 h at 4°C and ethanol was decanted before samples were resolved in 6 M urea/ 2 M thiourea buffer. Finally, samples were reduced, alkylated, digested and desalted as described above (phosphoproteomics).

### NanoLC-MS/MS analysis

Phosphopeptides and peptides from whole cell lysates were separated on a MonoCap C18 High Resolution 2000 column (GL Sciences) at a flow rate of 300 nl/min. 6 h and 4 h gradients were performed for whole cell peptides and phosphopeptides, respectively. Peptides from HA-affinity purified samples were separated on 45 min, 2 h or 4 h gradients with a 250 nl/min flow rate on a 15 cm column (inner diameter: 75 μm), packed in-house with ReproSil-Pur C18-AQ material (Dr. Maisch GmbH). A Q Exactive Plus instrument (Thermo Fisher) was operated in the data-dependent mode with a full scan in the Orbitrap followed by 10 MS/MS scans, using higher-energy collision dissociation. For whole proteome analyses, the full scans were performed with a resolution of 70,000, a target value of 3×10^6^ ions, a maximum injection time of 20 ms and a 2 m/z isolation window. The MS/MS scans were performed with a 17,500 resolution, a 1×10^6^ target value and a 60 ms maximum injection time. For phosphoproteome analysis, the full scans were performed with a resolution of 70,000, a target value of 3×10^6^ ions and a maximum injection time of 120 ms. The MS/MS scans were performed with a 35,000 resolution, a 5×10^5^ target value, 160 ms maximum injection time and a 2 m/z isolation window. For AP-MS of M97 interactomes, full scans were performed at a resolution of 70,000, a target value of 1×10^6^ and maximum injection time of 120 ms. MS/MS scans were performed with a resolution of 17,500, a target value of 1×10^5^ and a maximum injection time of 60 ms. Isolation window was set to 4.0 m/z. A Q-Exactive HF-X instrument (Thermo Fisher) or Q-Exactive Plus instrument was used for AP-MS samples of HHV kinases. The Q-Exactive HF-X instrument was run in Top20 data-dependent mode. Full scans were performed at a resolution of 60,000, a target value of 3×10^6^ and maximum injection time of 10 ms. MS/MS scans were performed with a resolution of 15,000, a target value of 1×10^5^ and a maximum injection time of 22 ms. Isolation window was set to 1.3 m/z. The Q-Exactive Plus instrument was run in data-dependent top10 mode. Full scans were performed at a resolution of 70,000 a target value of 3×10^6^ and maximum injection time of 120 ms. MS/MS scans were performed with a resolution of 35,000, a target value of 5×10^5^ and a maximum injection time of 120 ms. The isolation window was set to 4.0 m/z. In all cases normalized collision energy was 26.

### Data analysis

Raw data were analyzed and processed using MaxQuant 1.5.2.8 (M97 interactomes) or 1.6.0.1 (phosphoproteomics, whole cell lysates and interactomes of HHV-CHPKs) software (Cox and Mann, 2008). Search parameters included two missed cleavage sites, fixed cysteine carbamidomethyl modification, and variable modifications including methionine oxidation, N-terminal protein acetylation, asparagine/glutamine deamidation. In addition, serine, threonine, and tyrosine phosphorylations were searched as variable modifications for phosphoproteome analysis. Arg10 and Lys8 and Arg6 and Lys4 were set as labels where appropriate. The peptide mass tolerance was 6 ppm for MS scans and 20 ppm for MS/MS scans. The “match between runs” option was disabled and “re-quantify”,“iBAQ” (intensity based absolute quantification) and “second peptide” options were enabled. Database search was performed using Andromeda search engine (Cox et al., 2011) against a protein database of MCMV strain Smith and a Uniprot database of mus musculus proteins with common contaminants. Raw-data from AP-MS samples of HEK293-T cells were searched against a Uniprot Database of human proteins and the sequences of transgenic HHV kinases including common contaminants. False discovery rate was set to 1% at PSM, protein and modification site level. MS raw data and MaxQuant output tables have been deposited to the ProteomeXchange Consortium via the PRIDE partner repository (Perez-Riverol et al., 2019) with the dataset identifier PXD016334.

### Bioinformatics of phosphoproteomic profiles

Phosphosite data and whole proteome data were filtered to exclude contaminants, reverse hits and proteins only identified by site (that is, only identified by a modified peptide). Phosphorylation sites were ranked according to their phosphorylation localization probabilities (P) as class I (P > 0.75), class II (0.75 > P > 0.5) and class III sites (P < 0.5). Class I sites (in at least one of the replicates) were used with a multiplicity of one (that is, only one phosphorylation site on a peptide). MaxQuant normalized site ratios (from Phospho (STY)Sites.txt file) were used and corrected by the ratio of the corresponding protein (from proteinGroups.txt file) for the respective replicate. SILAC ratios of replicates were log2 transformed, averaged and sites were considered that were quantified in at least one of the replicates. Sites were then categorized as belonging to nuclear or cytosolic proteins, based on the GO annotation of the source protein. Source proteins and corresponding sites with no clear nuclear or cytosolic annotation were classified as “no category”. To assess differences in subcellular phosphoproteomic profiles we used the average SILAC ratio of cells infected with RXLmut and LXFmut virus and compared phosphosites that belong to nuclear or cytosolic proteins (see above). One-sided wilcoxon-rank sum test were performed comparing these two subsets of phosphosites with any possible amino acid in the region +4 to −4 (0 refers to the phosphorylated amino acid). Comparisons were considered when there were at least 19 phosphosites from both cytosolic and nuclear proteins for an amino acid at a given position quantified.

### Bioinformatics of HHV-CHPK interactomes

AP-MS data were filtered as described above, ratios were log2 transformed and replicates were averaged (mean) when they were quantified in all three replicates. Two-sided one sample t-tests (null hypothesis: µ_0_=0) were performed on the experimental data and a set of simulated data where enrichment ratios were permuted for the individual replicates (999 permutations). The t-test p-values were then adjusted according to the permuted data. The p-values in Table S1, Figure 1 and Figure S1 were adjusted in this way. Candidate interactors were selected based on a combination of adjusted p-value and means of the three replicates. To harmonize the data obtained from the different CHPK-IPs, we discriminated candidate interactors from background binders based on volcano plots. For all IPs, we used a fixed p-value cut-off of 0.05 and a flexible SILAC fold-change cut-off according to a false discovery rate estimation. For this, we used the simulated data as a false positive set and accepted candidate interactors above a SILAC fold-change that recalled maximum 1 % false positives.

FDR calculations were based on the simulated data as false positives. This yielded a set of high-confidence candidate interactors for IPs with individual HHV kinases. To compare individual prey proteins across the IPs with different kinases we imputed missing values with random values from a normal distribution (with mean 0 and standard deviation 0.25). Enrichment profiles were clustered using Euclidean distance and assembled into a heatmap using R. GO enrichment of the prey proteins in selected sets of clusters were performed using Metascape tool (Tripathi et al., 2015).

### Bioinformatics of M97 interactomes

AP-MS data were filtered as described above, ratios were log2 transformed and replicates were averaged (mean) when they were quantified in at least two of the replicates. Two-sided one sample t-tests (null hypothesis: µ_0_=0) were performed on the experimental data and proteins were considered as interactor when they were below a t-test p-value of 0.05 and above a log2 SILAC fold-change of 1.5. Additionally, the molar amount of bait and prey proteins in immunoprecipitates was estimated by iBAQ values. Therefore, the iBAQ values were summed up for samples where M97 was precipitated, sorted and log10 transformed.

### Immunoblot analysis

Whole cell lysates were prepared and adjusted to equal protein concentrations as described previously (Zydek et al., 2010). Subcellular fractionation into nuclei and cytoplasmic extracts was performed as described (Méndez and Stillman, 2000). Proteins were resolved by sodium dodecyl sulfate (SDS) polyacrylamide gel electrophoresis (PAGE) and blotted to polyvinylidene fluoride membranes, following standard procedures (Sambrook and Russell, 2001). To prevent non-specific binding, blots were incubated in Tris-buffered saline/ 0.1% Tween-20 (TTBS)/ 5% skim milk. Afterwards blots were incubated with the following primary antibodies: Cyclin A, C19 (Santa Cruz); cyclin E, clone M20 (Santa Cruz); CDK2, clone M2 (Santa Cruz); HA, clone 3F10 (Roche); M99, mouse antiserum (generously provided by Khanh Le-Trilling, University Hospital Essen); IE1, clone Croma 101; M45, clone M45.01; M57, clone M57.02 (all obtained from Center for Proteomics, Rijeka). The blots were developed using horseradish peroxidase-conjugated secondary antibodies in conjunction with the *Super Signal West Dura* chemiluminescence detection system (Thermo Fisher).

### Immunoprecipitation

Cells were lysed by freezing-thawing in immunoprecipitation buffer (IPB): 50 mM Tris-Cl pH 7.4, 150 mM NaCl, 10 mM MgCl_2_, 10 mM NaF, 0.5 mM Na_3_VO_4_, 0.5% Nonidet P-40, 10% glycerol, 1 mM dithiothreitol (DTT), 2 µg/ml aprotinin, 1 mM leupeptin, 1 mM Pefabloc. Cell extracts were clarified by centrifugation at 20.000 g. Cyclin A immunoprecipitations were performed as previously described (Eifler et al., 2014). For HA immunoprecipitations, a µMACS HA isolation kit (Miltenyi Biotec) was employed according to the manufacturer’s instructions, except that IPB was used as both lysis and washing buffer.

### Kinase assay

First, HA-M97 was immunoprecipitated from infected cells. To this end, IPB extracts were prepared and incubated with HA antibody clone 3F10 (Roche) and Protein G-conjugated agarose beads. The immunoprecipitates were washed several times with IPB and twice with 20 mM Tris-Cl (pH 7.4), 10 mM MgCl_2_, 1 mM DTT. Then, precipitates were incubated under constant agitation for 60 min at 30°C in kinase reaction buffer: 20 mM Tris-Cl (pH 7.4), 10 mM MgCl_2_, 1 mM DTT, 10 mM β-glycerophosphate, 50 μM ATP, 5 μCi of [γ-^32^P]ATP. Kinase reactions were analyzed by 8% SDS-PAGE followed by autoradiography.

### Immunofluorescence microscopy

3T3 fibroblasts were grown and infected on glass coverslips. Where indicated, cells were incubated with 10 μM 5-ethynyl-2′-deoxyuridine (EdU) for 30 min. To analyze for M97 localization and sites of EdU incorporation, coverslips were washed with PBS and incubated for 10 min in 4% paraformaldehyde/PBS fixation solution, followed by additional washing and incubation in PBS-T permeabilization solution (PBS, 0.1% Triton X-100, 0.05% Tween 20) and 2% bovine serum albumin (BSA) fraction V/PBS-T blocking solution. Afterwards, incorporated EdU was conjugated to Alexa Fluor 488 using the Click-iT EdU labeling kit (Thermo Fisher). Then samples were incubated overnight at 4°C with the following antibodies: anti-HA clone 3F10 (Roche) or anti-M57 clone M57.02 (Center for Proteomics, Rijeka), both diluted to 1 μg/ml in 2% BSA/PBS-T. After washing in PBS, cells were incubated for 1 h at 25°C with Alexa Fluor 488 or 647-coupled anti-IgG antibodies (Thermo Fisher). Coverslips were mounted on glass slides in 4′,6-diamidino-2-phenylindole (DAPI) containing Fluoromount-G medium (Thermo Fisher). Images were acquired by an Eclipse A1 laser-scanning microscope, using NIS-Elements software (Nikon Instruments). Equal microscope settings and exposure times were used to allow direct comparison between samples.

### Flow cytometry

Flow cytometry of DNA content and viral IE gene expression was performed as described (Zydek et al., 2010). Anti-IE1 (clone Croma 101, CapRI) and anti-M57 (clone m57.02, CapRI) were used as primary, Alexa 647 conjugated goat anti-mouse IgG (Thermo Fisher) as secondary antibody reagents.

## ACKNOWLEDGMENTS

The authors thank Jens von Einem, Tihana Lenac and Khan-Le Trilling for sharing valuable reagents. This work was supported by a grant (#900005) from the Joachim-Herz-Stiftung to L.W. M.Sc. was funded by a MD student research stipend from the Berlin Institute of Health.

## AUTHOR CONTRIBUTION

Conceptualization, B.B., L.W.; Methodology, B.B., M.Sc., H.W., K.I., E.O., M.Se., L.W.; Formal analysis, B.B., H.W., E.O.; Investigation, B.B., M.Sc., H.W., I.G., K.I., E.O., L.W.; Resources, M.Sc., I.G., B.V., L.W.; Data curation, B.B.; Writing (Original Draft), B.B., M.Sc., L.W.; Funding Acquisition, M.Sc., W.B., M.Se., C.H., L.W.; Supervision, W.B., M.Se., C.H., L.W.

## SUPPLEMENTAL INFORMATION

**Figure S1 (related to Figure 1).**
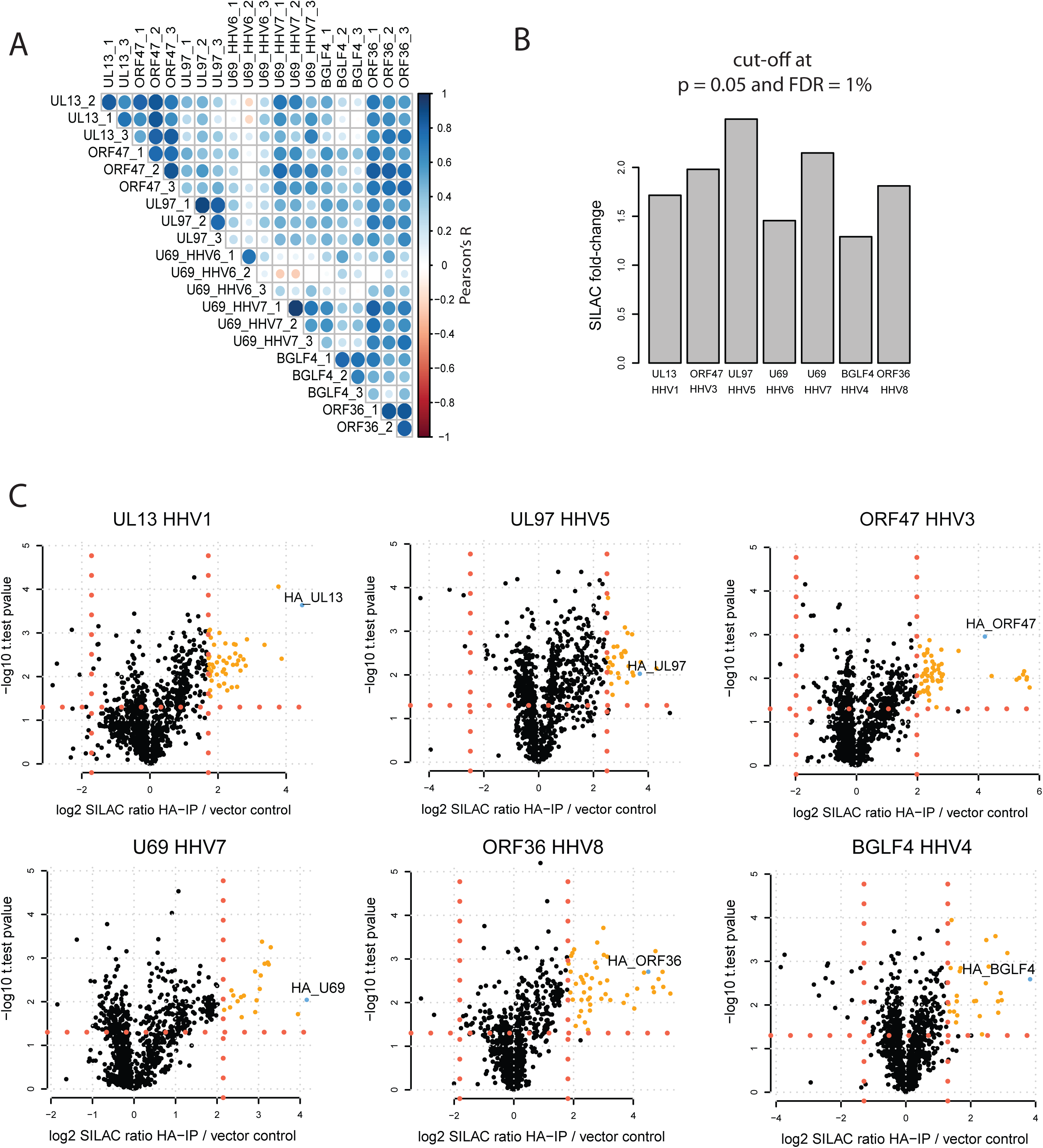
Data analysis of the protein-protein Interactome of HHV kinases. **(A)** Pearson’s correlation coefficients (Pearson’s R) comparing individual AP-MS experiments and replicates. **(B)** SILAC fold-change cut-offs when a t-test p-value of 0.05 and a FDR at 1% was used. **(C)** Individual volcano plots for IPs with six different herpesviral kinases. Candidate interaction partners above the fold-change and p-value cut-offs (indicated by red dotted lines) are highlighted orange.

**Figure S2 (related to Figure 1).**
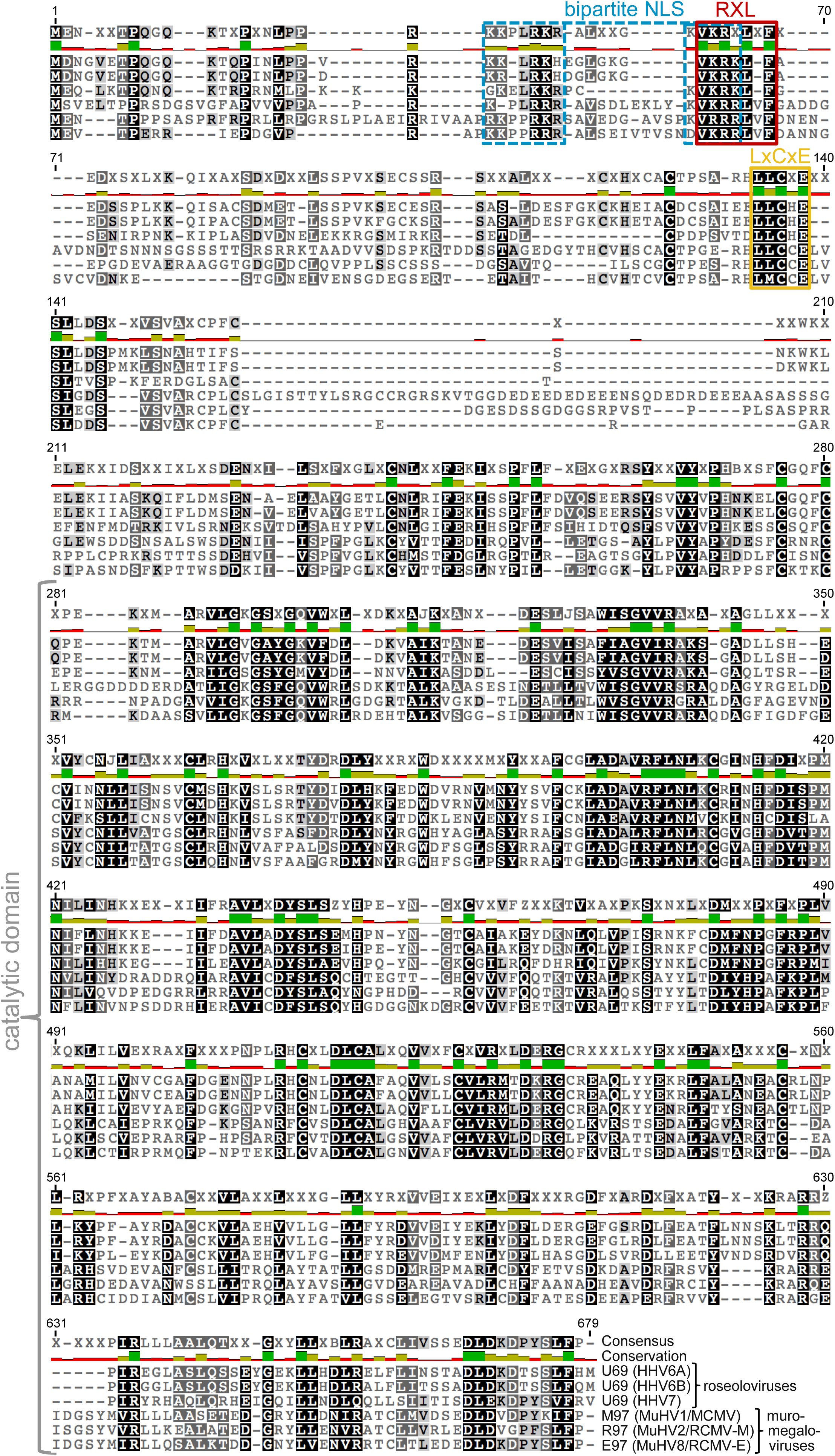
Multiple sequence alignment of β-herpesviral kinases. **(A)** Multiple sequence alignment was performed including kinases of roseolo- and muromegaloviruses. The conserved catalytic domain is indicated by brackets. The conserved bipartite NLS (blue), the RXL/Cy motif (red) and LXCXE motif (yellow) localize to the otherwise less-conserved N-terminal domain.

**Figure S3 (related to Figure 3).**
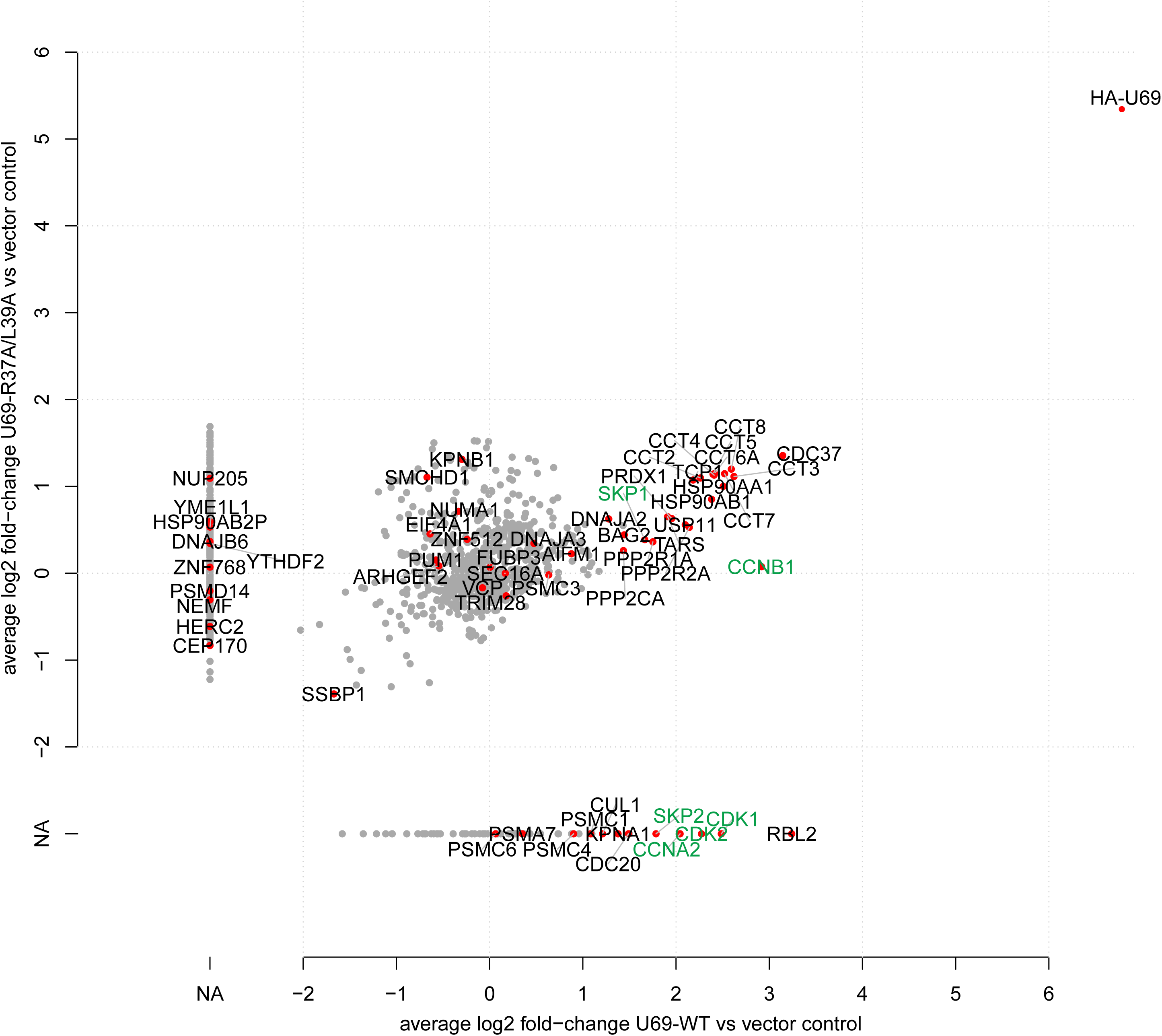
The RXL/Cy-motif assembles higher-order Cyclin-CDK complexes. SILAC-labeled HEK293-T cells were transfected with WT or R37A/L39A mutant versions of HHV6-U69. 24 h post transfection samples were subjected to AP-MS and shotgun proteomics. The enrichment of proteins precipitating with WT versus vector control was plotted against the enrichment of mutant versus vector control SILAC log2 fold-changes. Gene names are given for proteins that are interacting with any of the herpesviral kinases (Supplementary Table 1). Cyclins, CDKs or associated factors are highlighted green.

**Figure S4 (related to Figure 3).**
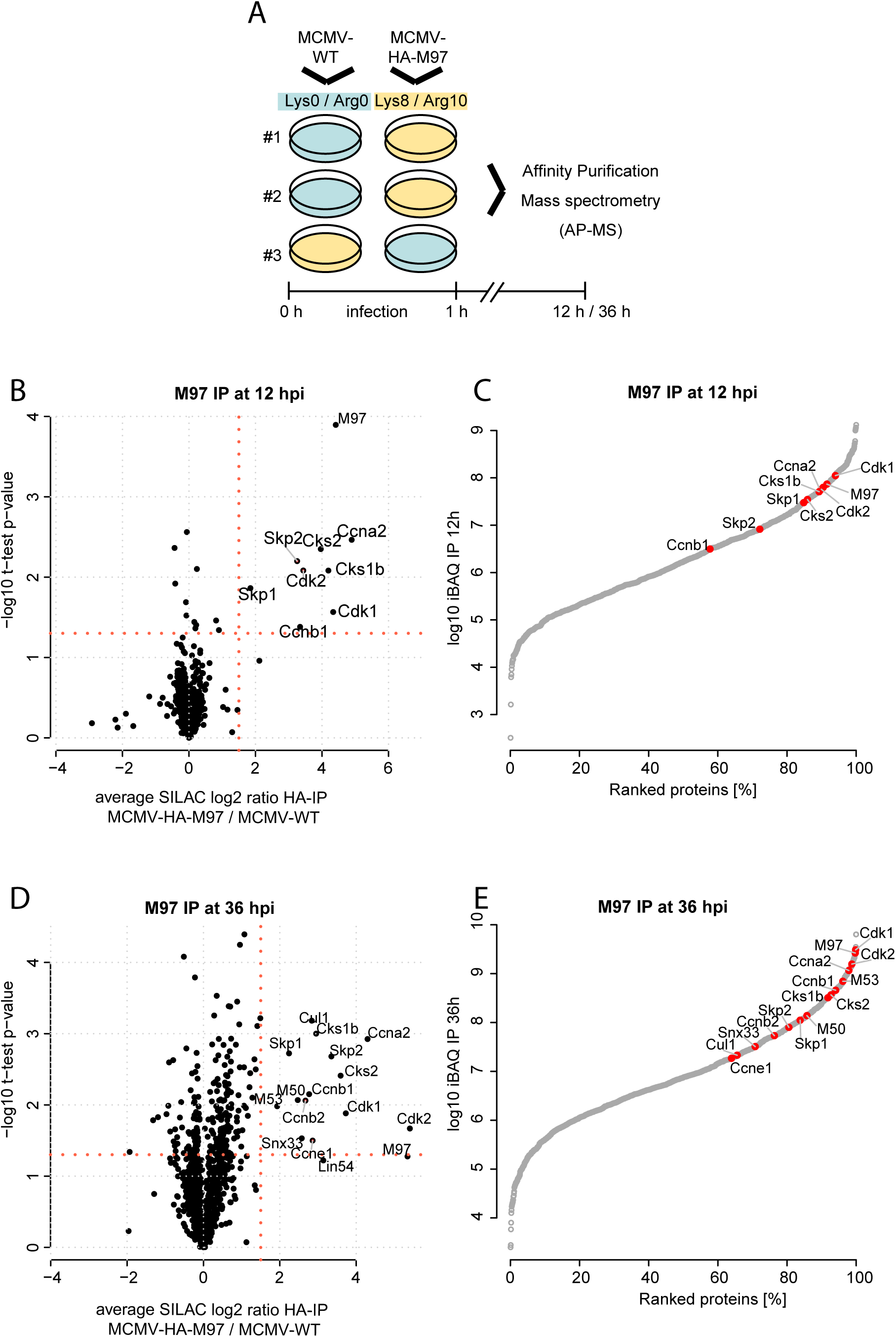
The time-resolved interactome of M97 during infection. **(A)** Experimental setup. SILAC Heavy and Light labeled cells were infected with MCMV-WT virus or a MCMV strain that expresses HA-tagged M97. Samples were subjected to affinity-purification mass-spectrometry. **(B, D)** The SILAC ratio MCMV-HA-M97 / MCMV-WT served to discriminate interactors of M97 from background binders. The average SILAC fold change and the t-test p-value of the three biological replicates is depicted for the IP at 12 **(B)** and 36 **(D)** hours post infection. **(C, E)** iBAQ values were calculated in HA-M97 precipitates at 12 **(C)** and 36 **(E)** hours post infection. Specific interactors at 12 and 36 hours post infection, respectively, are highlighted red. iBAQ values correlate with the molar amount of proteins in a sample and give thus estimates on the stoichiometry between individual protein complex members.

**Figure S5 (related to Figure 3).**
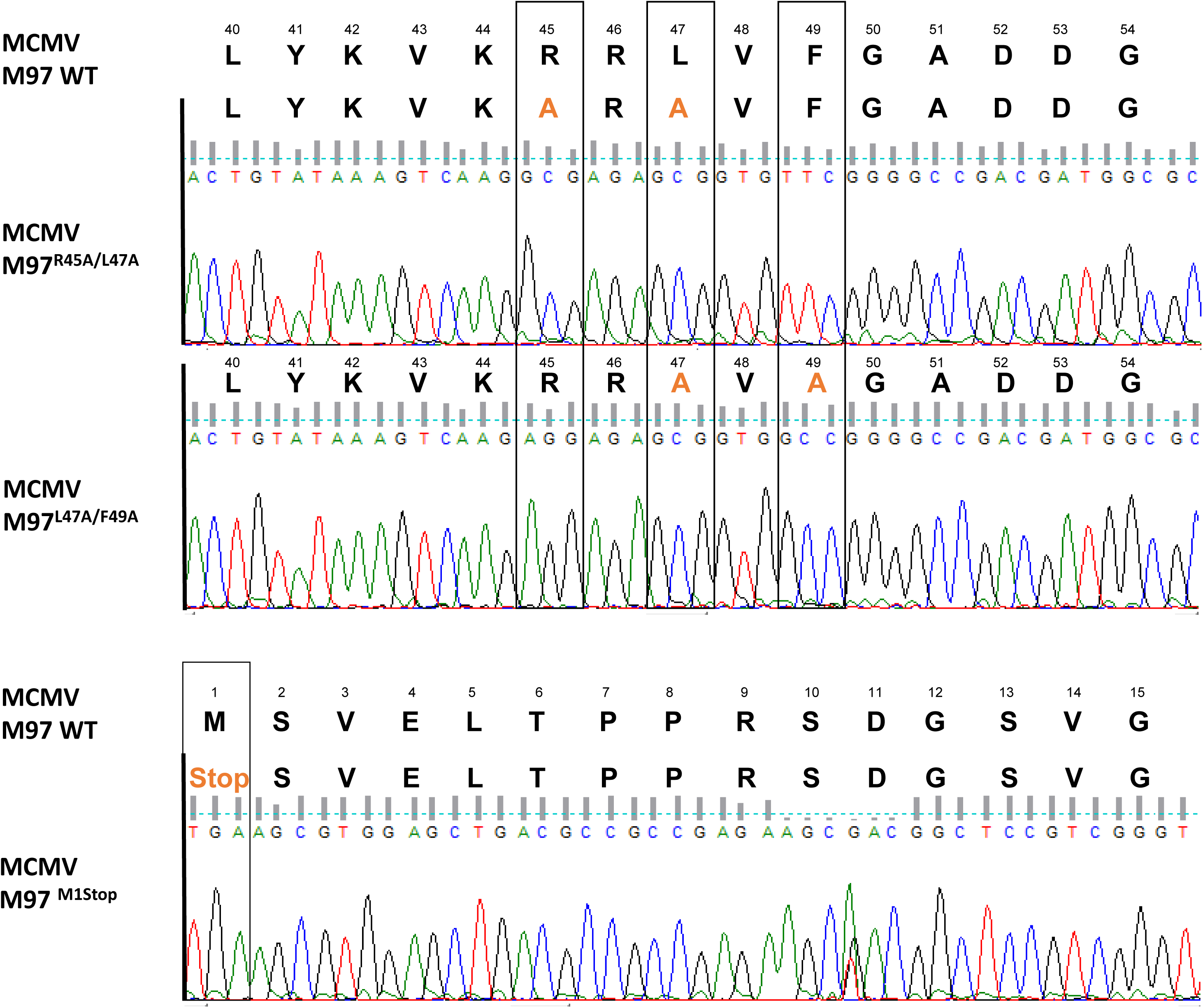
Sanger Sequencing of mutated MCMV-strains. DNA-sequence chromatograms resulting from Sanger sequencing of the mutated M97 gene region from the indicated recombinant viruses. The amino acids marked in orange highlight the introduced mutations.

**Figure S6 (related to Figure 4).**
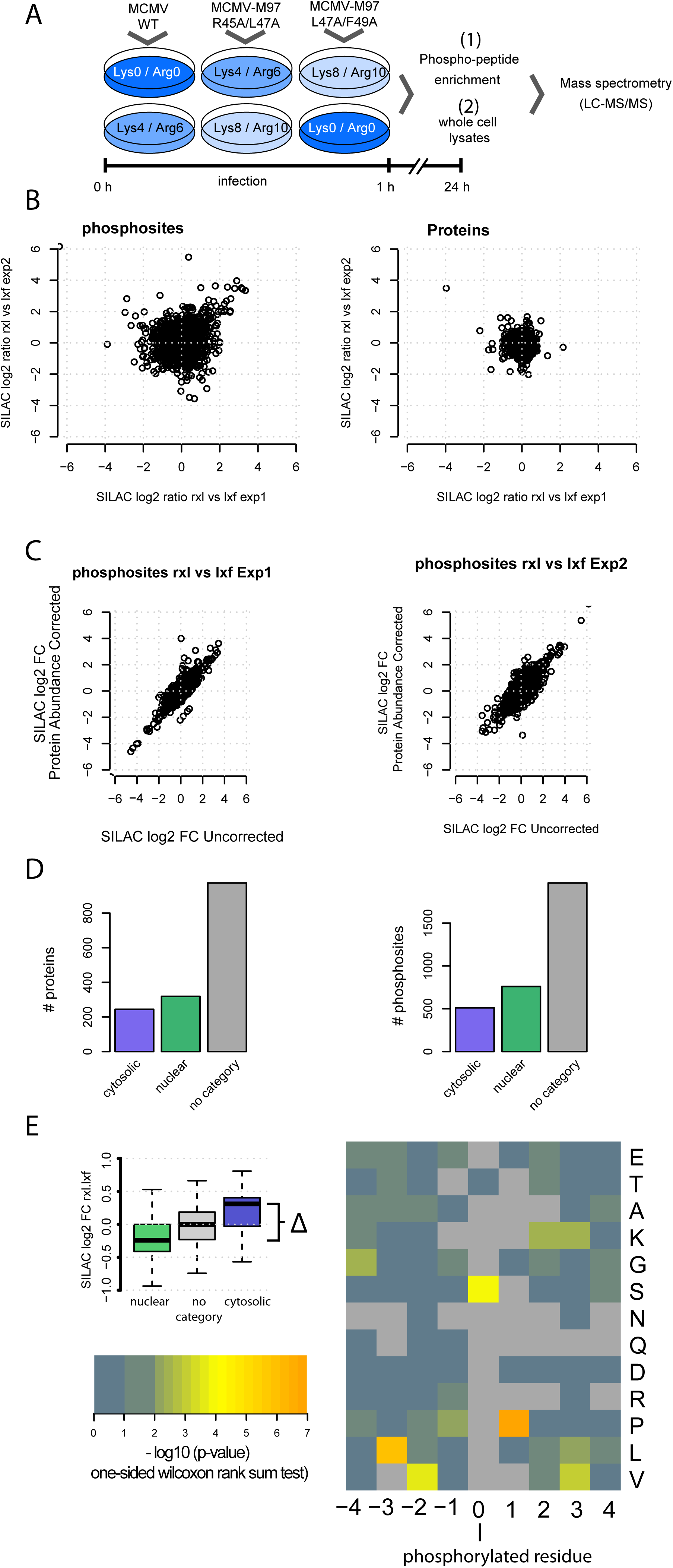
Differences in subcellular phosphoproteomic profiles upon infection with M97 mutant viruses. **(A)** Experimental setup. SILAC Heavy, Medium and Light labeled cells were infected with MCMV-WT, M97-R45A/L47A or M97-L47A/F49A viruses. At 24 hours post infection cells were harvested and either subjected to a phosphoproteomics workflow or whole proteome analysis. Note that only the direct SILAC ratio M97-R45A/L47A / M97-L47A/F49A was used for further analysis. **(B)** Quantified phosphosites and Proteins in both replicates in the SILAC comparison M97-R45A/L47A versus M97-L47A/F49A. **(C)** Quantified phosphosites were corrected by the SILAC fold-change of the respective source protein for both replicates. **(D)** Quantified proteins (left panel) and phosphosites (right panel) were classified as cytosolic, nuclear or “no category” (that is, no clear nuclear or cytosolic GO annotation). The number of proteins or phosphosites that were classified is given. **(E)** Global comparison of phosphosites on nuclear proteins with phosphosites on cytosolic proteins by screening any possible amino acid flanking the phosphorylated residue. Log_10_ P values (wilcoxon rank sum test) for the differences in nuclear and cytosolic phosphosites of each amino acid in the region −4 to +4 flanking serine or threonine (position: 0). Consistent with existing data on pUL97 (Oberstein et al., 2015), we found significant differences for phosphorylated Serines, followed by Prolines at position +1, Lysines at position +2 and +3 as well as hydrophobic residues at positions −2/-3/+3.

**Figure S7 (related to Figure 5).**
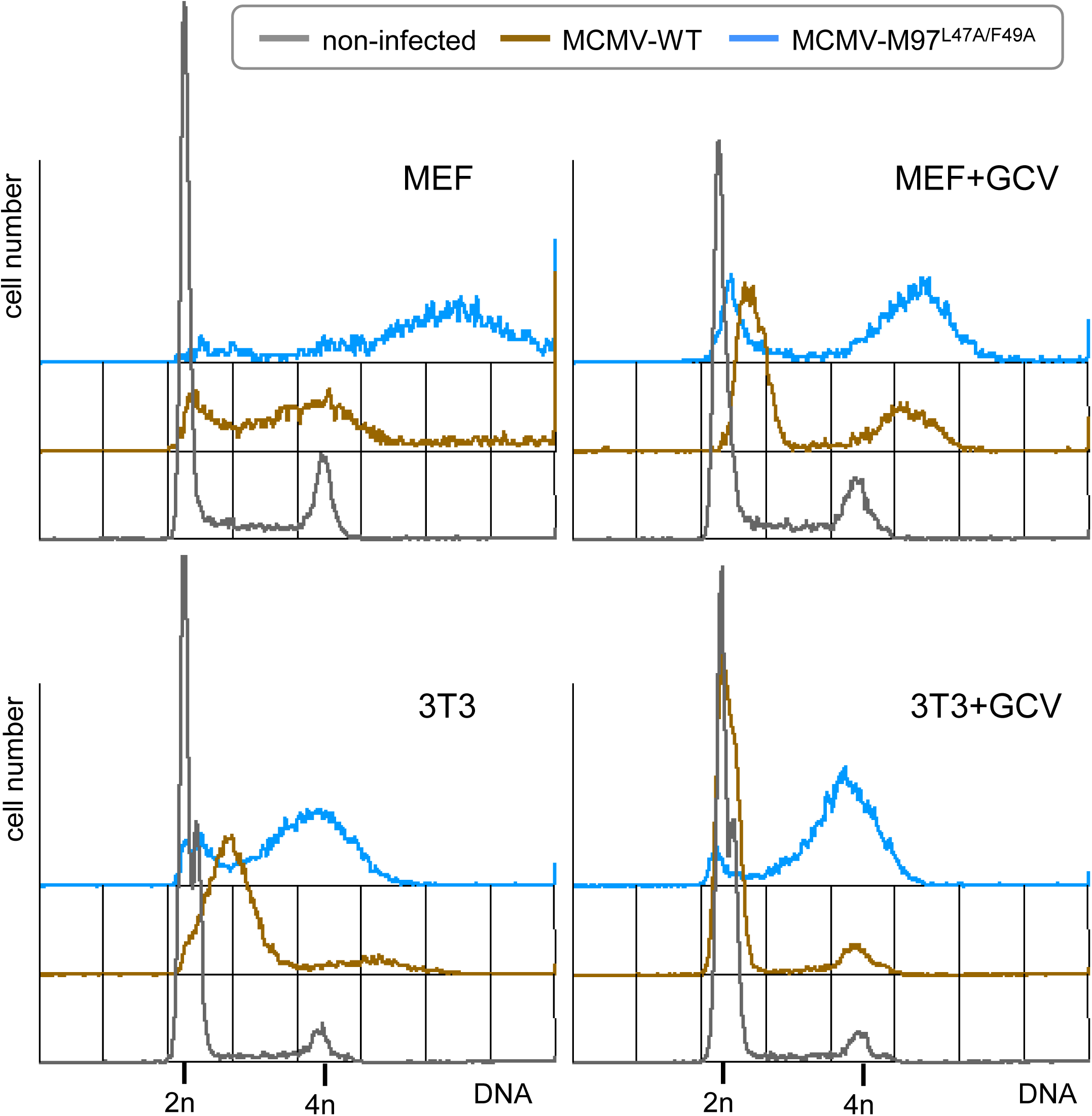
Loss of M97-Cyclin A interaction leads to unscheduled cellular DNA synthesis and over-replication in primary cells. Mouse embryonic fibroblasts (MEF) or NIH-3T3 fibroblasts were synchronized in G0/G1 phase by growth factor deprivation (0.05% serum for 2 days). Cells were infected with M97-RXL/Cy mutant MCMV or the parental WT virus (MOI=2). After removal of the virus inoculum, low serum conditions were maintained. Where indicated, 50 μM ganciclovir (GCV) was added to the culture medium. At 48 h post infection, cells were harvested and analyzed by flow cytometry for DNA content and viral infection markers (IE1 for GCV-treated cells, M57 for untreated cells). DNA histograms of infected and non-infected cell populations are shown.

